# FSCN1-Mediated Hepatic Gluconeogenesis Is Indispensable for Neonatal Mice Survival

**DOI:** 10.1101/2025.05.18.654691

**Authors:** Xiangxiang Liu, Yuanzhao Hu, Liangwei Wu, Yiwen Zhang, Lei Sang, Yake Gao, Lei He, Wenyong Xiong, Shengyu Yang, Jianwei Sun

## Abstract

Actin-Bundling Protein Fascin1 (FSCN1) is encoded by the *Fscn1* gene, and crucial for cytoskeletal remodeling and cellular migration. While previous study linked *Fscn1* deficiency to neonatal lethality in mice, the underlying metabolic mechanism remains unexplored. Here, in this study, we report that systemic knockout (KO) of *Fscn1* leads to 52.2% mortality within 24 hours post-birth, accompanied by severe hypoglycemia in KO pups compared to other littermates. Remarkably, this lethality was fully rescued by oral glucose administration, indicating a glucose supply-dependent survival mechanism. Surviving *Fscn1* KO neonates displayed persistent developmental deficits, including growth retardation and depleted lipid stores, despite intact canonical insulin-regulated hepatic gluconeogenic pathways. Transcriptomic profiling of P0 livers revealed that *Fscn1* loss predominantly disrupts metabolic pathway, within the glycerol phosphate shuttle being the most significantly downregulated module. Mechanistically, *Fscn1* KO livers exhibited markedly reduced protein levels of glycerol-3-phosphate dehydrogenase isoforms (GPD1/GPD2), key enzymes bridging glycolysis and gluconeogenesis. Consistently, glycerol tolerance tests (GTT) demonstrated impaired glycerol-to-glucose conversion in *Fscn1* KO mice, confirming defective glycerol-driven gluconeogenesis. Our findings establish FSCN1 as a novel cytoskeletal-metabolic integrator essential for neonatal survival by sustaining hepatic glucose production from glycerol, thus revealing an unexpected role of actin dynamics in coordinating metabolic adaptation during early postnatal development.

**Graphical Abstract:** 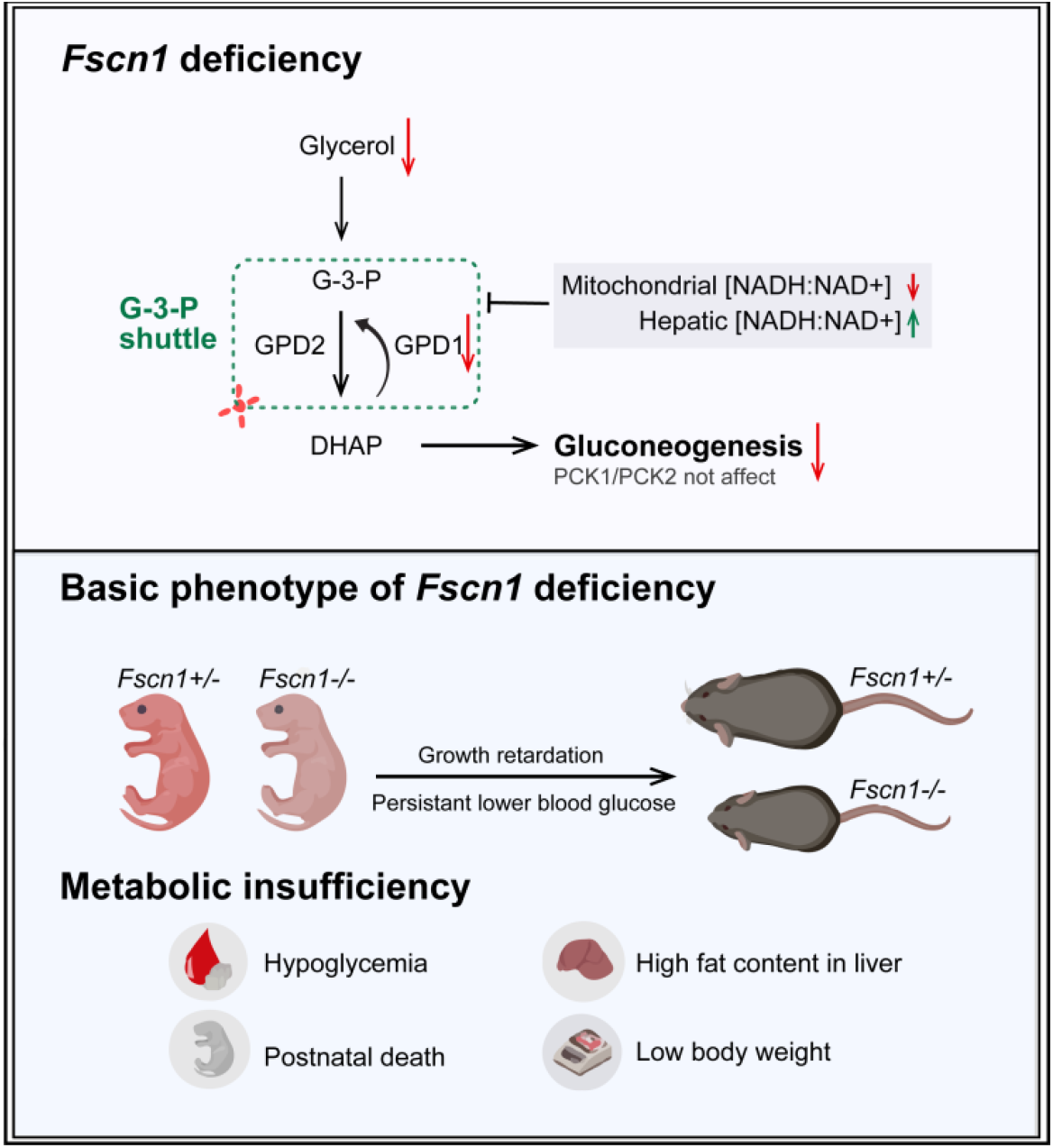

**Highlights:**

1. FSCN1 deficiency triggers hypoglycemia-driven neonatal mortality by disrupting hepatic glucose production.
2. *Fscn1* loss impairs the gluconeogenesis via suppressing GPD1/GPD2 expression, while sparing insulin-regulated gluconeogenesis.
3. Cytoskeletal dysfunction impairs glycerol-to-glucose conversion, revealing actin-metabolism crosstalk in postnatal adaptation.

## 1 Introduction

*Fscn1*, a widely studied pro-oncogenic gene in the past decade, encodes the actin-bundling protein Fascin1 (also named as Fascin). It physically crosslinks filamentous actin (F-actin) contributes to the formation of invasive protrusions that facilitate cell migration, and adhesion [1-4]. Beyond its classical roles, FSCN1 modulates the actin-binding proteins [5-7], interacts with microtubules [8], and integrates with the linker of nucleoskeleton and cytoskeleton complex [9]. While FSCN1’s oncogenic roles cancer progression and metabolic homeostasis are well-documented [10-19], its physiological functions in normal organ development remain poorly understood.

In adult tissues, FSCN1 is expressed at low levels in dendritic cells, neurons and vascular endothelial cells, whereas its expression peaks during embryonic development, particularly in the neural and mesenchymal system [1]. Although murine studies initially suggested FSCN1 is dispensable during development [20, 21], *Fscn1* deficiency causes neonatal lethality post-birth without affecting embryo viability [22]. Early hypotheses attributed this lethality to postnatal feeding or respiratory challenges [22], with later work implicating lactation defects and feeding abnormalities in *Fscn1* KO females [23]. However, our observation of impaired survival and persistent hypoglycemia in offspring derived from *Fscn1* KO points to intrinsic metabolic dysregulation independent of maternal factors. This study investigates the role of FSCN1 in hepatic metabolism during neonatal adaptation using a systemic *Fscn1* KO mouse model.

Hepatic gluconeogenesis—the synthesis of glucose from non-carbohydrate precursors such as glycerol, lactate, and amino acids — is tightly regulated by hormones, redox, and substrate availability [24, 25]. While adults rely on hormone-driven transcription of enzymes like glucose-6-phosphatase (G6PC), and phosphoenolpyruvate carboxykinase 1 (PCK1) [26, 27]. Neonates prioritize glyceroneogenesis, a glycerol-centric pathway critical for adapting to lipid-rich maternal milk. This pathway begins with glycerol phosphorylation by glycerol kinase (GK), followed by conversion to dihydroxyacetone phosphate (DHAP) via glycerol-3-phosphate dehydrogenase (GPD1/GPD2), ultimately fueling gluconeogenesis [28]. Neonates undergo rapid metabolic shifts post-birth, marked by the elevated glucagon and decreased insulin levels, which rapidly activate hepatic gluconeogenesis to meet the brain’s high glucose demand [29]. Importantly, neonatal Gluconeogenesis contributes 8-9% of total glucose production-a threshold indispensable for survival [30]. Despite evidence linking *Fscn1* deficiency to neonatal lethality via hypoglycemia, the mechanistic basis of this metabolic dysregulation remains unresolved. This study investigates whether impaired gluconeogenic capacity underlies the hypoglycemia-driven mortality observed in *Fscn1*-deficient neonates.

Herein we demonstrate that FSCN1 is indispensable for neonatal hepatic gluconeogenesis. Postnatal *Fscn1* KO mice showed severe metabolic dysregulation, characterized by persistent hypoglycemia and impaired glycerol-driven gluconeogenesis. The lethality in *Fscn1* KO mice can be rescued through glucose supplements. These findings identify that FSCN1 orchestrates metabolic reprogramming during neonatal adaptation response, establishing its essential role in maintaining hepatic glucose production and ensuring newborn survival.

## 2 Materials and Methods

### 2.1 Animals

All animal experiments were conducted in accordance with protocols approved by the Institutional Animal Care and Use Committee of the Yunnan University (Ethical Committee Approval No. SINH-2020-DQR-3). Wild-type C57BL/6J mice were obtained from the Animal Center of Yunnan University, *Fscn1* knockout mouse model was generated with deletion of 543 bp across 2 exons and maintained under a specific pathogen-free (SPF) condition.

Embryonic specimens were collected from timed-pregnant *Fscn1*^*+/-*^ dams at embryonic day 19 (E19) via uterine dissection. A heterozygous intercross breeding strategy (*Fscn1*^*+/-*^ ♀ × *Fscn1*^*+/-*^ ♂) was implemented to analyze Mendelian inheritance patterns, with neonatal genotypes determined by PCR amplification [31] (primers in Table S1) and quantified for distribution analysis. Postnatal day 0 (P0) littermates were utilized for dual-omics profiling (transcriptomics and metabolomics), while 6-week-old littermates were assessed for growth retardation phenotypes through longitudinal body weight measurements. Metabolic evaluations, including insulin tolerance tests (ITT) and glycerol gluconeogenesis assays (glycerol tolerance tests), were performed on P35 mice (see Table S2 for reagents). Age-matched (± 3 days) littermate pairs were prioritized for comparative analyses to control for environmental variability. Sex determination via *Sry* gene analysis [32] confirmed no sex-based disparities in glucose levels or ATP production (*p* >0.05). Wild-type (WT) controls were generated from an independent C57BL/6J breeding colony to ensure enough experimental animals, as the number of WT littermates from heterozygous crosses were less.

### 2.2 Cell culture and genetic manipulation

Mouse embryonic fibroblasts (MEFs), human embryonic kidney 293T (293T) cells, and human hepatocellular carcinoma cell line (Huh7) were acquired from the American Type Culture Collection (ATCC). All cell lines were maintained as adherent cultures in DMEM (Gibco, Shanghai, China), supplemented with 10% fetal bovine serum (ExCellBio, Suzhou, China) and 1% penicillin/streptomycin (Solarbio, Beijing, China) under standard conditions (37°C, 5% CO_2_) without Mycoplasma contamination. For genetic perturbation, shRNA constructs targeting *Fscn1* (sh*Fscn1*) were subcloned into the pLKO.1-TRC vector, while CRISPR/Cas9 guide RNAs (sg*Fscn1*) were engineered into the Lenti-CRISPR-V2 backbone. Oligonucleotide sequences for RNA interference and gene editing are cataloged in Table S2, with vector construction protocols adapted from established methodologies (https://www.addgene.org/protocols/plko/, https://www.zlab.bio/guide-design-resources). All expression vectors underwent Sanger sequencing verification (Tsingke, Beijing, China) prior to lentiviral packaging and cellular transduction.

### 2.3 Glucose measurement and metabolic challenge tests

Neonatal blood samples (P0) were collected in tubes with 2% EDTA-coated tubes (Biofil, Guangzhou, China) and centrifuged 3000 rpm for 10 mins at 4°C to isolate plasma. Quantitative glucose analysis was performed using a glucose oxidase-based assay (Nanjing Jiancheng Bio, Nanjing, China), and the detailed methods are as follows: 5 μL plasma aliquots were combined with 200 μL reaction reagent in 96-well plates (BioFil, Guangzhou, China), incubated at 37°C for 10 mins, and absorbance measured at 505 nm using Biotek microplate reader with ddH_2_O serving as negative control. For fasting studies, age-matched WT, *Fscn1*^*+/-*^ and *Fscn1* KO mice underwent controlled fasting (16 hours: 18:00-10:00) with ad libitum access to water, followed by paired fed/fasted blood sampling via tail vein puncture. Glycerol tolerance test (glycerol-mediated gluconeogenesis test) were conducted in P35 mice from *Fscn1*^*+/-*^ and *Fscn1*^*-/-*^ groups following 16 hours overnight fasting. After baseline glucose measurement, mice were numbered, weighed, and received oral gavage of 2.5% glycerol solution (2.73 mM in saline; Sangon Biotech, Shanghai, China) at 10 mL/kg body weight, based on referenced methods for glycerol quantification [27, 33]. Serial glucose measurements were monitored at 0, 30, 60, 90, and 120 minutes post-administration using standardized tail nick procedures. Area under the curve (AUC) was calculated using trapezoidal method. All experimental protocols were synchronized with facility light cycles (06:00-18:00) to minimize circadian variability.

### 2.4 Multi-Omics profiling

Neonatal littermates (*Fscn1*^*+/-*^ *vs Fscn1*^*-/-*^) were genotyped within 24 hours postpartum via tail biopsy PCR prior to hepatic tissue collection. Liver specimens (n=3-4/per group) were homogenized in TriZol reagent (Lablead, Beijing, China), and stored at -80°C until RNA extraction. Paired-end RNA sequencing libraries were prepared and constructed in Novogene (Beijing, China) with a read length of 2 × 150 bp on an Illumina NovaSeq 6000 platform. Both raw data and clean data underwent FastQC v0.11.9 quality control, and index construction was performed with Bowtie2 (http://bowtie-bio.sourceforge.net/bowtie2/index.shtml), while genome alignment of the clean data was conducted using TopHat2 (https://ccb.jhu.edu/software/tophat/index.shtml). Transcriptome assembly was carried out using Cufflinks (https://cole-trapnell-lab.github.io/cufflinks/). Gene expression levels were calculated based on the fragments per kilobase million reads (FPKM) method. A t-test was utilized to calculate p-values for differentially expressed genes (DEGs) with | log2 (foldchange) | ≥ 1, considering p-values <0.05 as significant (DEGs). The raw transcriptome data were obtained, and subsequent analysis was performed as previously described [34].

Untargeted metabolomics analysis employed liquid nitrogen-snap-frozen P0 hepatic samples (n=3, per group) processed through cryogenic milling, methanol: acetonitrile: water (2:2:1) extraction and lyophilization, which were strictly followed the Agilent metabolomics sample preparation procedure [35]. Reconstituted extracts were analyzed on an Agilent 6545 Q-TOF LC/MS system equipped with Agilent SB C18 column (2.1×100 mm, 1.8 μm) using gradient elution, with data acquisition in positive/negative ESI modes and solvent blanks as procedural controls.

### 2.5 Immunoblotting

Liver tissues were dissected and immediately homogenized in ice-cold RIPA lysis buffer (Beyotime Biotechnology, Nanjing, China) containing protease/phosphatase inhibitors cocktail (MCE, Shanghai, China). After 1-hour incubation on ice, lysates were centrifuged at 12,000 rpm for 15 minutes at 4°C. Protein quantification were performed using a BCA kit (Thermo, Shanghai, China) with bovine serum albumin standards. After protein denaturation with the loading buffer, 10-12.5% gradient SDS-polyacrylamide gels (Epizyme Biotech, Shanghai, China) were employed for immunoblotting experiments. Following wet transfer, the PVDF membranes (Millipore, Shanghai, China) were blocked with 5% non-fat milk in TBS-T (Tris-buffered saline with 0.1% Tween-20) for 1 hour at room temperature, followed by overnight incubation at 4°C with primary antibodies: anti-GPD1 (ABclonal, Wuhan, China) and anti-GPD2 (Proteintech, Wuhan, China). After three TBST washes, membranes were incubated with HRP-conjugated secondary antibodies, for 1 hour at 4°C. The proteins were visualized using the ECL-plus western blot detection system (Tanon-5200 Multi). The primary antibody was diluted to 1:1000, and secondary antibody was diluted to 1:10000. Additionally, detailed information regarding the antibodies used can be found in Table S3.

### 2.6 RNA isolation and real-time quantitative PCR

Total RNA extracted from hepatic tissues and cell lines using TriZol Reagent (Lablead, Beijing, China). RNA purity and concentration were determined spectrophotometrically (A260/A280 ratio 1.8-2.0, NanoDrop 2000). Reverse transcription was performed with 1 μg RNA using the First-strand cDNA Synthesis Mix (Lablead, Beijing, China). Quantitative PCR amplifications were conducted in triplicate using 2x Realab Green PCR Fast mixture (Lablead, Beijing, China) on a Real-Time PCR System (Bio-rad) under the following cycling conditions: initial denaturation at 95°C for 2 minutes, 40 cycles of 95°C for 10 s and 60°C for 30 s, with melt curve analysis (65-95°C, 0.5°C/s increments). Gene expression quantification employed the 2^-ΔΔCt^ method [36]. The sequences of primers used are listed in Table S2.

### 2.7 Pathological examination and immunity-related experiments

Hepatic specimens were immediately fixed in 4% paraformaldehyde (PFA) in phosphate-buffered saline (pH 7.4) for 24 hours at 4°C. Fixed tissues underwent graded ethanol dehydration (70%-100%), xylene clearing, and paraffin embedding. Serial 5-μm sections were cut with a rotary microtome (Leica) and mounted on poly-L-lysine-coated slides (Citotest, Nanjing, China). Hematoxylin and eosin (H&E) staining was performed according to previously established procedures [37]. Specifically, H&E staining solution was applied for 2 minutes, followed by differentiation using an acidic differentiation solution for 5 seconds, and the sections were washed for 10 minutes to achieve a blue coloration. After a series of ethanol dehydration and xylene clearing steps, 2-3 drops of neutral gum were immediately added to the sections, which were then sealed and dried at 37°C.

### 2.8 Measurement of lipids-related components

Serum and hepatic lipid profiles were comprehensively analyzed in in WT, *Fscn1*^*+/-*^, and *Fscn1*^*-/-*^ mice at P0 and P35. Serum samples were centrifuged at 3000 rpm for 15 minutes at 4°C, while liver homogenates were prepared through sequential centrifugation at 12000 rpm for 15 minutes twice at 4°C. The levels of non-esterified fatty acids (NEFA) in serum and liver tissues were quantified using an automatic biochemical analyzer, following the manufacturer’s instructions. Glycerol levels were assessed using a glycerol assay, with lipid components including triglycerides (TG), total cholesterol (T-CHO), high-density lipoprotein cholesterol (HDL-C), and very low-density lipoproteins cholesterol (VLDL-c) levels measured under standardized conditions. Hepatic ketone bodies β-hydroxybutyrate (BOH) and acetoacetate (AcAc) were analyzed using a ketone assay kit. All metabolite concentrations were normalized to total protein content by BCA assay, and experimental details for commercial kits are cataloged in Table S2.

### 2.9 Insulin level and Insulin tolerance test

To investigate the effect of insulin signaling on blood glucose levels, both insulin level and insulin tolerance test (ITT) were conducted. Due to limited serum volume from P0 neonates, systemic insulin levels were evaluated in P35 mice (*Fscn1*^*+/-*^ *vs Fscn1*^*-/-*^) following 4-hour fasting (08:00-12:00). Blood was collected via retro-orbital puncture into serum separator tubes. Insulin concentrations were determined using a mouse-specific ELISA kit (Jonln Biotechnology, Shanghai, China). For the insulin tolerance test, age-matched mice (5-week-old) were fasted for 4 hours (08:00-12:00), with free water access, and then weighed. After baseline glucose measurement, mice were intraperitoneally injected with recombinant human insulin (Novo Nordisk, Copenhagen, Denmark) at 0.5 U/kg body weight. Serial glucose measurements were performed via tail vein sampling at 0, 15, 30, 60, and 120 minutes post-injection.

### 2.10 Statistical analysis

Experimental cohorts were randomized from standardized breeding litters. No statistical methods were employed to predetermine sample sizes. Compared with *Fscn1*^*+/-*^ mice, 462 genes with |log2FoldChange|≥ 1 and P value < 0.05 were screened by using R software limma package for differential analysis [38]. Quantitative densitometry of western blot bands was performed using Image J software. All datasets were analyzed in GraphPad Prism 9.0.1, presented as mean ± SEM (standard error of mean) from n≥3 independent biological replicates. Intergroup comparisons utilized two-tailed unpaired Student’s t-tests, with p-values less than 0.05 considered significant. Survival curves were generated through Kaplan-Meier methodology.

## 3 Results

### 3.1 *Fscn1* deficiency triggers hypoglycemia-induced neonatal lethality

To investigate FSCN1’s physiological role, we generated systemic *Fscn1* knockout (KO) mice model (Fig. 1A). Genotyping confirmed a 543 bp deletion spanning two *Fscn1* exons, PCR amplification using the primer pairs *Fscn1-genome-*F and *Fscn1-genome-*R produces a single band in WT mice. whereas heterozygous (*Fscn1*^*+/-*^) exhibits dual amplicons (1620 bp and 1077 bp) corresponding to WT and *Fscn1* KO mice (Fig. 1B). *Fscn1* KO mice were identified using the deletion-specific primer pair *Fscn1-del* and *Fscn1-genome-*R, which produces a single band (739 bp) in WT and heterozygous mice, but no band in homozygous *Fscn1* KO mice (Fig. 1C). Western blotting analysis further confirmed the absence of FSCN1 in *Fscn1* KO mice (Fig. 1D).

**Figure 1.**
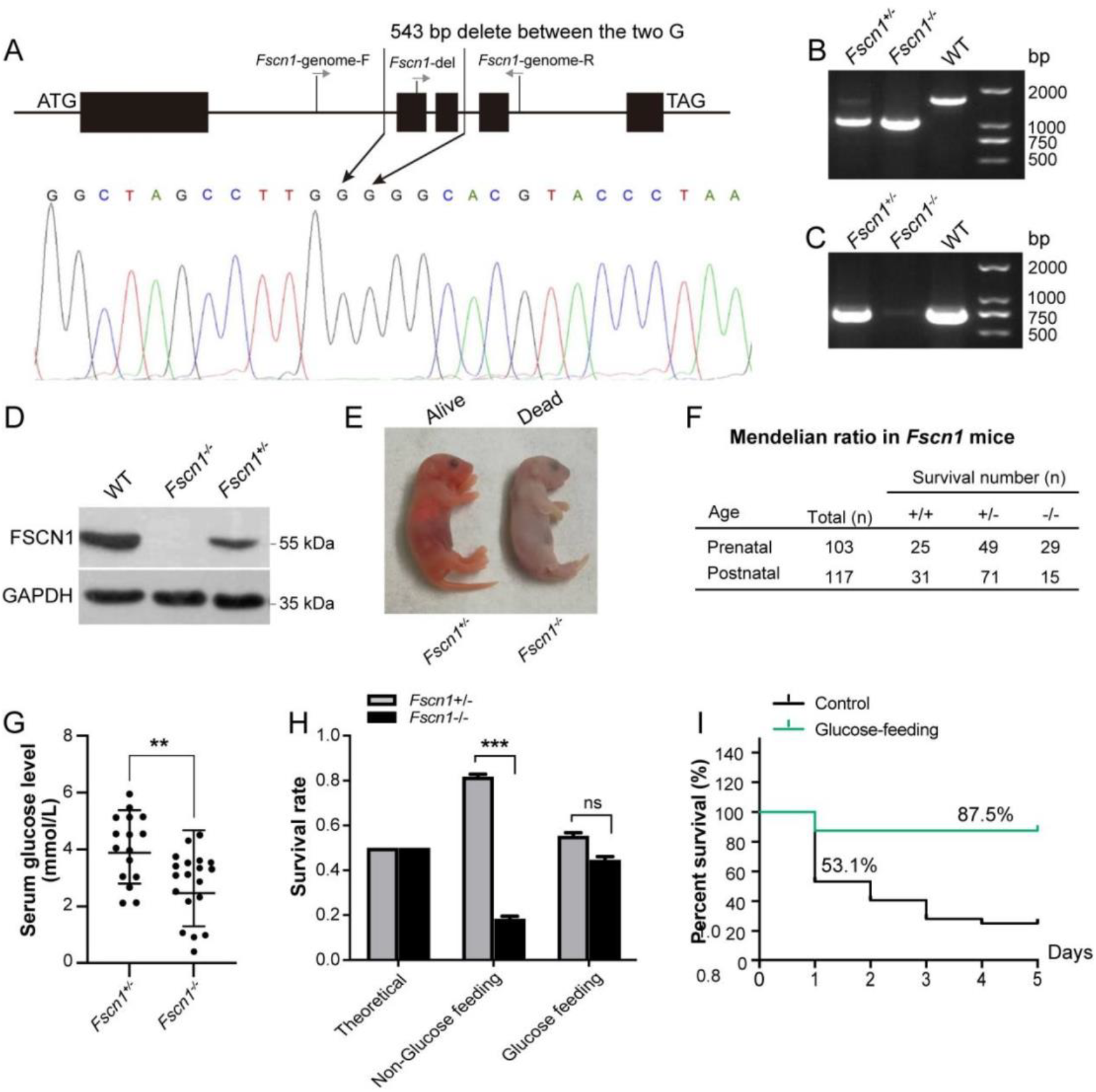
Systematic knockout of *Fscn1* induces hypoglycemia and high neonatal mortality. (A) Schematic diagram illustrating the construction of a systemic *Fscn1* knockout. (B) PCR analysis confirming wild-type (WT) mice, indicated by a 1620 bp band, using primers *Fscn1-genome-*F and *Fscn1-genome-*R. (C) PCR analysis identifying *Fscn1*^*-/-*^ mice, characterized by the absence of 739 bp band, using primers *Fscn1-del* and *Fscn1-genome-*R. (D) Western blotting verification of the absence of FSCN*1* protein level in liver tissue of knockout mice. (E) Representative image showing *Fscn1* knockout neonates that died within 24 hours after birth. (F) Statistical analysis of *Fscn1*^*+/-*^ offspring ratios obtained via heterozygous mating (♀ *Fscn1*^*+/-*^ *&* ♂ *Fscn1*^*+/-*^). (G) Serum blood glucose levels in *Fscn1*^*+/-*^ and *Fscn1*^*-/-*^ P0 mice. (n=16 for *Fscn1*^*+/-*^; n=19 for *Fscn1*^*-/-*^). (H) Offspring survival ratio of *Fscn1*^*+/-*^ and *Fscn1*^*-/-*^ P0 mice after oral glucose supplementation within 24 hours post-birth (Glucose content is 100 g/L). Lethality rate of non-glucose feeding *Fscn1*^*-/-*^ mice: 47.8% (ratio, 11/23), lethality rate of Glucose feeding *Fscn1*^*-/-*^ mice: 20% (ratio, 4/20). (I) Survival curves of *Fscn1*^*+/-*^ and *Fscn1*^*-/-*^ mice (P0) within the first 5 days postnatally, with or without oral glucose supplementation. (n=32 for control; n=16 for glucose feeding group). Mice used in these experiments were littermates. Data are presented as mean ± SEM (standard error of mean), statistical significance was determined by two-tailed t-test, \**p* < 0.05, \*\*\**p* < 0.001.

Strikingly, *Fscn1*^*-/-*^ neonates exhibited postnatal lethality within 24 hours post-birth (Fig. 1E). To further investigate these findings, we employed a mating strategy involving female/male heterozygotes (♀ *Fscn1*^*+/-*^ *&* ♂ *Fscn1*^*+/-*^). Upon examination of plug in the fertilized female, we focused on mice in the embryonic period of E18.0 (Prenatal) and within 24 hours of birth (Postnatal). Genotypic analysis of prenatal mice demonstrated mice from heterozygous offspring nearly adhered to the Mendelian inheritance ratio 1:2:1 (WT : *Fscn1*^*+/-*^ : *Fscn1*^*-/-*^ = 25 : 49 : 29) (Fig. 1F). Notably, 47.8% of *Fscn1*^*-/-*^ neonates died postnatally during the first 24 hours of life (Fig. S1A), indicating that *Fscn1* deficiency induced neonatal death predominantly occurred in the postnatal period within 24 hours, which consistent with previous reports [22].

Given the importance of energy supply for neonatal survival, we measured serum and liver glucose levels. *Fscn1* KO neonates exhibited severe hypoglycemia, with significantly reduced glucose concentrations in both serum (3.76 ± 1.47 mM *vs*. 5.57 ± 0.77 mM in *Fscn1*^*+/-*^ littermates) and liver homogenate (Fig. 1G, S1B). Consistent with AMPK’s role in glucose sensing [39], *Fscn1* KO neonates showed increased hepatic AMPK phosphorylation at Thr192, a hallmark of energy stress, accompanied by decreased ATP levels (Fig. S1C-D), indicating energy deprivation due to *Fscn1* deficiency. Remarkably, oral glucose supplementation (100 g/L) rescued survival rates from 53.1% to 87.5% (Fig. 1H-I), directly linking neonatal lethality to glucose insufficiency. and the mortality of *Fscn1*^*-/-*^ P0 mice decreased from 47.8% to 20% after glucose supplementation (Fig. S1A). These findings identify hypoglycemia as the primary driver of neonatal lethality in *Fscn1*-deficient mice, establishing glucose insufficiency as the central metabolic defects in this model.

### 3.2 Postnatal hypoglycemia drives developmental defects independent of insulin signaling

To evaluate the long-term effects of *Fscn1* deficiency, we longitudinally monitored body weights in untreated littermates from postnatal day 14 (P14) to P42. To minimize neonatal stress, mice were identified via toe marking at P14. *Fscn1* KO mice exhibited pronounced growth impairment, with significantly reduced body size and persistent weight deficits (P14-P42), compared to *Fscn1*^*+/-*^ littermates (Fig. 2A-B). Given the association between dysregulated cholesterol metabolism and growth retardation [40, 41], we examined lipid levels in P35 mice. Surprisingly, *Fscn1* KO mice displayed systemic hypolipidemia, characterized by reductions in total cholesterol (T-CHO), high-density lipoproteins (HDL-C), non-esterified fatty acid (NEFA) and triglycerides (TG), relative to WT and *Fscn1*^*+/-*^ counterparts (Fig. 2C). This broad lipid depletion suggests a global metabolic dysfunction underlying the growth phenotype. Notably, *Fscn1* KO mice maintained chronically depressed blood glucose levels into adulthood (6 weeks old, Fig. 2D), indicating a developmental failure to establish glucose homeostasis. The persistence of hypoglycemia beyond the neonatal period underscores the essential role of FSCN1 in metabolic programming during early development and warrants further investigation of insulin signaling dynamics.

**Figure 2.**
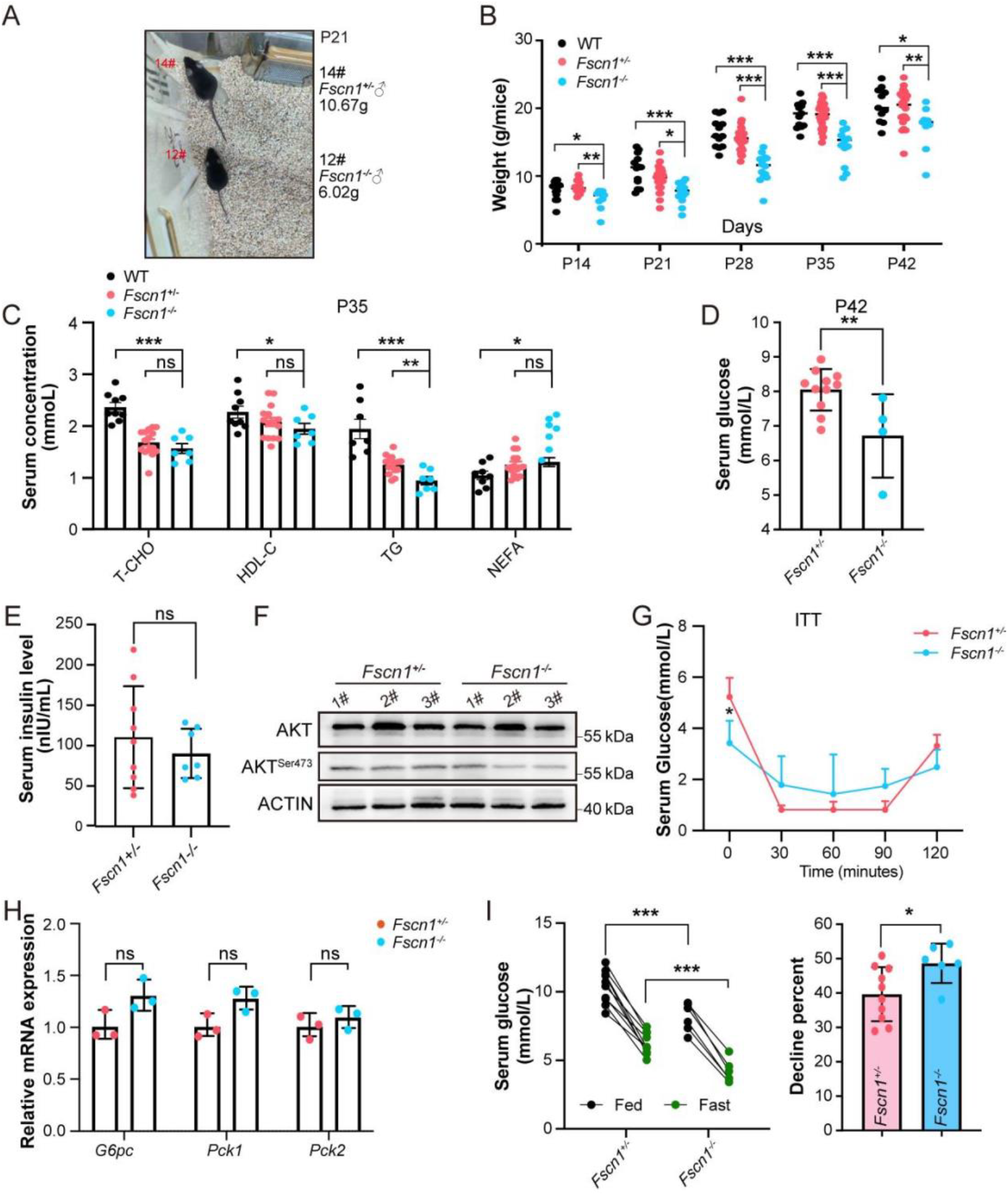
*Fscn1* deficiency induces persistent hypoglycemia and growth retardation in mice. (A) Images depicting body size differences between *Fscn1*^*+/-*^ and *Fscn1*^*-/-*^ mice at postnatal day 21 (P21), highlighting differences in overall growth. (B) Body weight measurements of *Fscn1*^+/-^ and *Fscn*1^-/-^ mice from 2 to 6 weeks (each group, n≥9). (C) Lipid profile analysis in P35 mice, including total cholesterol (T-CHO), high-density lipoprotein cholesterol (HDL-C), triglycerides (TG) and non-esterified fatty acid (NEFA) (each group, n≥7, except for NEFA measurements, which were performed on 3 mice per group). (D) Serum glucose levels in *Fscn1*^*+/-*^ and *Fscn1*^*-/-*^ P42 mice (*Fscn1*^*+/-*^, n=11; *Fscn1*^*-/-*^, n=4). (E) Serum insulin concentrations in P35 mice (*Fscn1*^*+/-*^, n=9; *Fscn1*^*-/-*^, n=7). (F). Western blot analysis of AKT phosphorylation in *Fscn1*^*+/-*^ and *Fscn1*^*-/-*^ P0 liver (each group, n=3). (G) Insulin tolerance test in P35 mice administered 0.5 U/kg insulin after 4 hours of fasting, (each group, n≥6). (H) Relative mRNA levels of key gluconeogenesis-related genes (*G6pc, Pck1* and *Pck2*) in the livers of *Fscn1*^*+/-*^ and *Fscn1*^*-/-*^ mice at P0 (each group, n=3). (I) Effects of starvation on blood glucose utilization in P35 mice (each group, n≥6). All mice used in these experiments were littermates, except WT mice. Data are presented as mean ± SEM (standard error of mean), statistical significance was determined by two-tailed t-test, *\*p* < 0.05, *\*\*p* < 0.01, *\*\*\*p* < 0.001.

Hyperinsulinemic hypoglycemia (HH), typically marked by high insulin secretion during hypoglycemia [42], was excluded as a mechanistic contributor in *Fscn1* KO mice. Serum insulin levels in KO mice remained comparable to *Fscn1*^*+/-*^ mice (Fig. 2E), with no evidence of enhanced insulin signaling, as evidenced by unaltered AKT phosphorylation at Ser473 (Fig. 2F). Insulin tolerance test (ITT) revealed mild insulin resistance in KO mice (Fig. 2G), likely a secondary consequence of chronic hypoglycemia rather than a primary driver of metabolic dysfunction. Critically, hepatic expression of canonical gluconeogenic regulators (*G6pc, Pck1, Pck2*) remained unaffected in *Fscn1* KO mice (Fig. 2H). Following a 16-hour fast, *Fscn1* KO mice exhibited a pronounced increase in glucose utilization (48.67% reduction *vs*. 39.68% in *Fscn1*^*+/-*^ controls; Fig. 2H), consistent with preserved glucose absorption but diminished reserves due to baseline hypoglycemia. Collectively, these findings exclude hyperinsulinemic and canonical gluconeogenic defects, instead implicating alternative pathways in *Fscn1* deficiency-associated hypoglycemia and growth impairment.

### 3.3 Hepatic transcriptomics reveals *Fscn1* is involved in metabolic regulation and glycerol-3-phosphate shuttle

While *Fscn1* deficiency does not impair canonical hormone-mediated gluconeogenesis, the underlying mechanisms of hypoglycemia in KO mice remained unresolved. Immunoblotting analysis revealed progressive postnatal decline of hepatic FSCN1 (Fig. S2A), highlighting its developmental stage-specific role in neonatal metabolic adaptation. To elucidate compensatory pathways, we conducted whole-liver transcriptome profiling of *Fscn1*^*+/-*^ and *Fscn1*^*-/-*^ neonates at P0. Principal component analysis (PCA) of the normalized RNA-seq data revealed clear genotype segregation, with principal components (PC1 and PC2) accounting for 61.8% and 12.7% of the total variance, respectively (Fig. 3B). A total of 462 differentially expressed genes (DEGs) were identified, with 235 genes up-regulated and 227 genes down-regulated in *Fscn1* KO mice, compared to *Fscn1*^*+/-*^ mice (Fig. 3C, 3G). Enrichment analysis of the DEGs highlighted significant Gene Ontology (GO) terms, including gluconeogenesis, nucleotide binding, and membrane dynamics (Fig. 3D-E). Kyoto encyclopedia of genes and genomes (KEGG) pathway analysis revealed 18 significantly perturbed pathways (p < 0.05), including metabolic pathways, chemical carcinogenesis-receptor activation, and steroid hormone biosynthesis (Fig. 3F). Of particular interest were *Gpd1* and *Gpd2*, encoding rate-limiting enzymes of the glycerol-3-phosphate (G-3-P) shuttle — a critical system channeling cytosolic glycerol into mitochondrial gluconeogenic flux (Fig. 3G) [33, 43, 44]. Transcriptional downregulation of *Gpd1* and *Gpd2* in *Fscn1* KO neonates suggest defective glycerol-to-glucose conversion, a key adaptation enabling the neonatal transition from placental nutrition to autonomous metabolic regulation.

**Figure 3.**
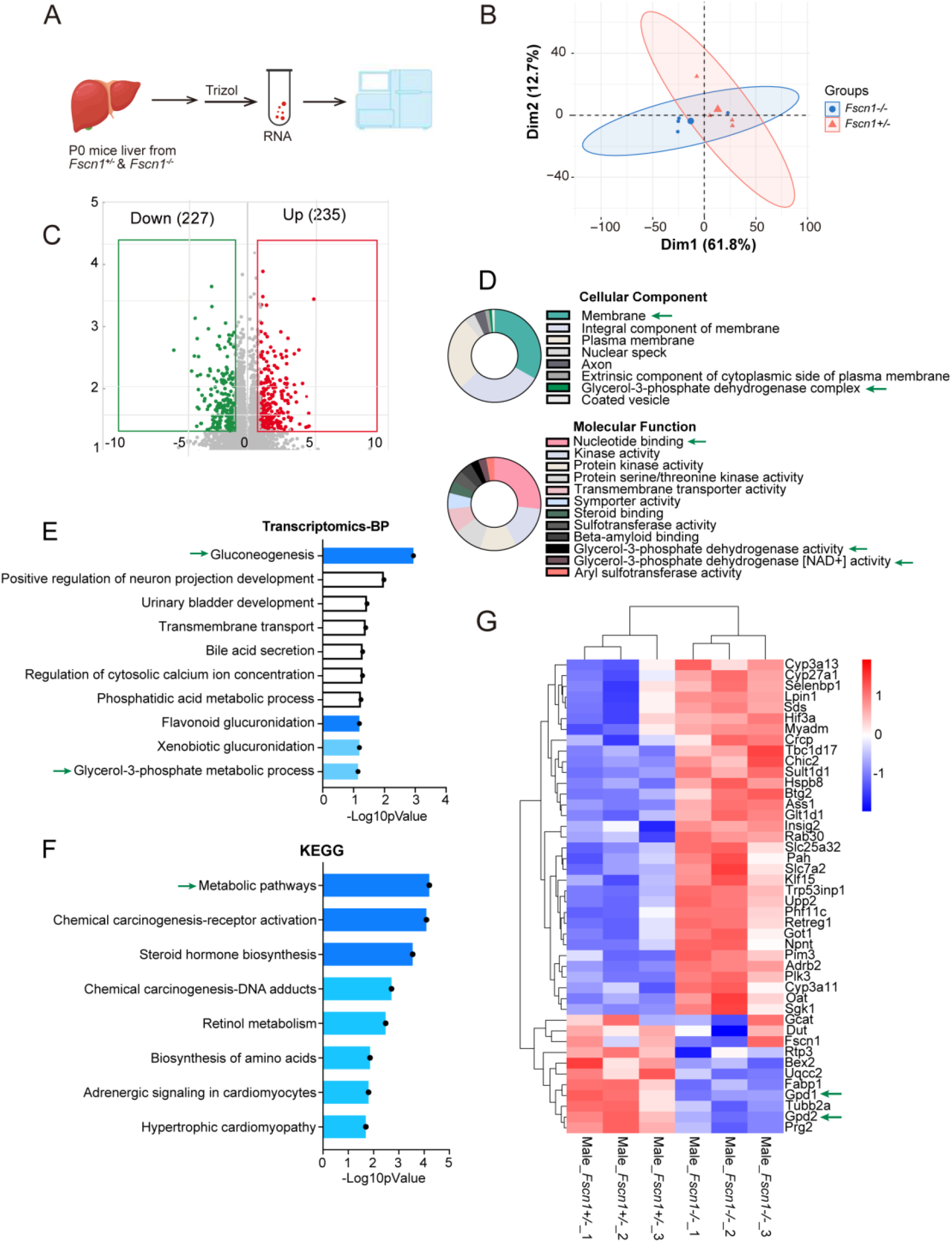
*Fscn1* deficiency affected the glycerol-3-phosphate process as revealed by transcriptomics analysis. (A) Schematic diagram illustrating the sample preparation process for transcriptomic sequencing. (B) Principal component analysis (PCA) of differentially expressed genes (DEGs). (C) Volcano plot depicting enriched DEGs identified from transcriptomic data. (D) Gene Ontology (GO) analysis of DEGs, categorized into cellular components (CC) and molecular functions (MF). (E) Significance analysis of biological process (BP) associated with DEGs. (F) KEGG pathway enrichment analysis of DEGs in the liver of *Fscn1*-deficient mice. (G) Heatmap illustrating of DEGs in the liver of *Fscn1*^*+/-*^ and *Fscn1*^*-/-*^ P0 mice. Mice used in above experiments were littermates. Data are shown as mean ± SEM, statistical significance was determined by two-tailed t-test, with *\*p* < 0.05, *\*\*\*p* < 0.001,

### 3.4 *Fscn1* regulates glycerol-driven hepatic gluconeogenesis via GPD1/2 -medicated Glycerol-3-Phosphate shuttle

To investigate FSCN1’s role in gluconeogenesis, we focused on GPD1 and GPD2, two essential enzymes controlling the glycerol-3-phosphate (G-3-P) shuttle, which links glycerol availability to mitochondrial redox balance [33]. *Fscn1* KO neonates exhibited significant reductions in both hepatic GPD1 and GPD2 protein and mRNA levels (Fig. 4A-B). To further elucidate FSCN1-dependent regulatory mechanism, we employed shRNA-mediated *Fscn1* knockdown and CRISPR-Cas9-mediated *Fscn1* knockout. shRNA targeting *Fscn1* in MEF specifically suppressed *Gpd1* transcripts without affecting *Gpd2* expression (Fig. 4C). Similarly, *Fscn1* KO in Huh7 cells recapitulated this specificity, showing reduced GPD1 but unchanged GPD2 (Fig. 4D). These models suggest that FSCN1 acts as a selective upstream regulator of GPD1, with GPD2 regulation appearing context-dependent,

**Figure 4.**
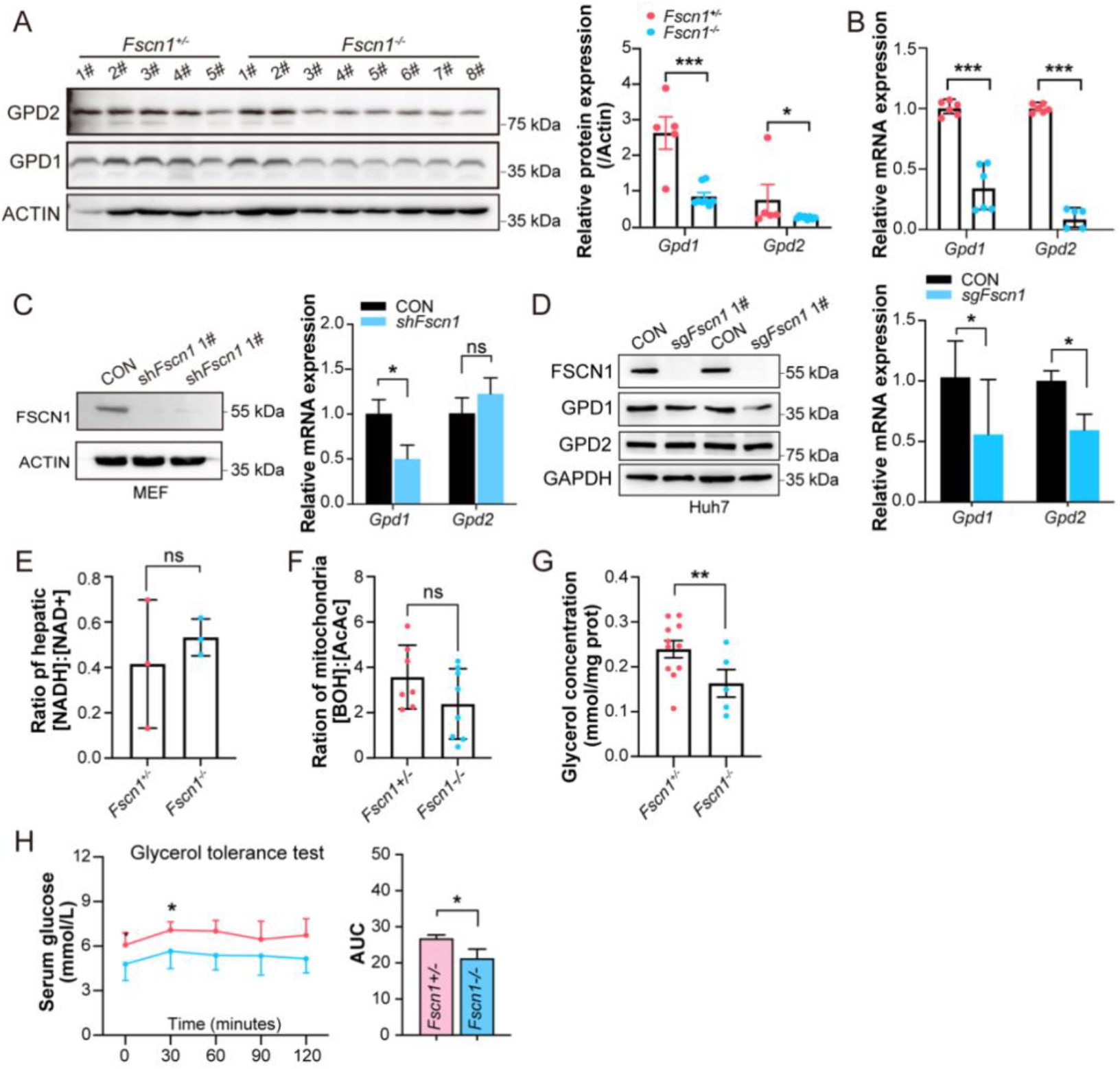
*Fscn1* deficiency impairs hepatic gluconeogenesis from glycerol. (A) Western blotting analysis of GPD1, GPD2 protein levels in P0 mice liver, with quantification using Image J (B) mRNA levels of *Gpd1, Gpd2* in P0 mice liver (n=3). (C) Validation of Fscn1, GPD1, GPD2 levels in Fscn1 knockdown MEF cells. (D) Protein and mRNA expression levels of GPD1 and GPD2 in sg*Fscn1*-Huh7 cells. (E) NADH/NAD^+^ ratio in the liver of *Fscn1*^*+/-*^ and *Fscn1*^*-/-*^ mice at P0 assessed by metabolomics. (F) Mitochondrial redox status measured by β-hydroxybutyrate (BOH) to acetoacetate (AcAc) ratio in P0 liver tissues. (G) Hepatic glycerol concentrations in *Fscn1*^*+/-*^ and *Fscn1*^*-/-*^ mice (P0). (H) Glycerol tolerance test in *Fscn1*^*+/-*^ and *Fscn1*^*-/-*^ P35 mice, with area under the curve (AUC) analysis. Mice fasted for 16 hours and administered 2.5% glycerol (n ≥ 6). All experiments used littermate mice. Data are presented as mean ± SEM, statistical significance was determined by two-tailed t-test, with, *\*p* < 0.05, *\*\*\*p* < 0.001.

To assess the functional consequences of GPD1 deficiency, we analyzed intracellular redox dynamics and substrate utilization. Metabolomic profiling revealed a significantly elevated [NADH:NAD^+^] ratio in *Fscn1* KO livers (Fig. 4E), indicating an elevated cytoplasmic redox state and preserved GPD1 enzymatic activity. Additionally, *Fscn1* KO livers accumulated very low-density lipoproteins (VLDL-c) and triglycerides (TG) (Fig. S2E). Since TG hydrolysis generates glycerol (for gluconeogenesis) and free fatty acids (for mitochondrial β-oxidation and ketogenesis). we evaluated mitochondrial function: *Fscn1* KO mice showed reduced ketone body levels (β-hydroxybutyrate [BOH] and acetoacetate [AcAc]; Fig. S2F), indicating impaired mitochondrial β-oxidation. Furthermore, the mitochondrial redox status was similarly disrupted in *Fscn1* KO mice (Fig. 4F). This coordinatedl dysregulation of cytoplasmic and mitochondrial redox homeostasis impairs the G-3-P shuttle’s capacity to transfer reducing equivalents, as evidenced by decreased hepatic glycerol levels in *Fscn1* KO mice (Fig. 4G).

To directly assess glycerol-driven gluconeogenesis, glycerol tolerance tests were performed in 5-week-old mice. *Fscn1* KO mice displayed blunted glycemic responses to oral glycerol administration (Fig. 4H), confirming a defect in glycerol-to-glucose conversion. Collectively, these findings demonstrate that *Fscn1* deficiency disrupts hepatic gluconeogenesis by impairing the GPD1-dependent G-3-P shuttle, ultimately leading to hypoglycemia and highlighting FSCN1’s essential role in neonatal survival and growth.

## 4 Discussion

Maintaining neonatal glucose homeostasis necessitates precise coordination between hepatic glucose production and peripheral utilization. In this study, we unveil an uncharacterized regulatory role for FSCN1 in regulating neonatal glucose metabolism, which is essential for neonatal survival. Unlike the canonical PCK1/PCK2-dependent gluconeogenesis pathway, FSCN1 deficiency induces hypoglycemia and growth retardation through disruption of the G-3-P shuttle bypassing canonical insulin-regulated gluconeogenic pathways. Importantly, *Fscn1* KO mice exhibited persistent hypoglycemia despite glucose supplementation rescuing neonatal lethality, accompanied by reduced body weight and lipid reserves. This metabolic defect occurs independently of hyperinsulinemia. Mechanistically, FSCN1 regulates GPD1-dependent gluconeogenesis, directly affecting the glycerol-to-glucose conversion process. Our findings identify FSCN1 as a novel metabolic regulator and highlight a critical link between cytoskeletal dynamics and hepatic glucose production, potentially providing insights into congenital hypoglycemia disorders associated with cytoskeletal dysfunction.

Neonatal hypoglycemia, affects approximately 10% of healthy term infants, predominantly within the first 24-48 hours post-birth [45, 46], attributable to high brain glucose demands and a large brain-to-body mass ratio [47]. While hypoglycemia can often be asymptomatic or present with nonspecific symptoms, severe cases (defined as blood glucose levels less than 40 mg/dL in term infants) can result in life-threatening complications including cardiac arrhythmias, seizures, and irreversible neurological damage [48, 49]. Prompt glucose supplementation is critical to prevent a cerebral energy crisis, due to the brain’s preferential reliance on glucose over ketones during acute hypoglycemia [50]. In our model, *Fscn1* KO neonates exhibited 52.2% lethality within 24 hours, indicating impaired postnatal metabolic adaptation. Clinical thresholds for hypoglycemia vary across etiologies [51], yet *Fscn1* KO neonates demonstrated significantly lower blood glucose levels (3.76 ± 1.47 mM) compare with their heterozygotes counterparts (5.57 ± 0.77 mM at P0), which is below the mammalian physiological threshold (∼5.0 mM) [52], aligning with diagnostic criteria of pathogenic hypoglycemia in human neonates [51]. The survival of *Fscn1* KO pups following oral glucose rescue underscores their glucose dependency, phenocopying starvation-induced hypoglycemia. This dependency is further evidenced by reduced hepatic ATP levels (1.05 ± 0.57 mM/L/mg prot) and compensatory AMPK activation in *Fscn1* KO pups, indicative of systemic energy deprivation. FFA, which undergo β-oxidation to generate ketone bodies for cerebral energy supply during fasting [29], also enhance gluconeogenic flux by providing acetyl-CoA and NADH for glucose synthesis [53, 54]. The concurrent reduction in ketone body levels in *Fscn1*-deficient neonatal, a direct metabolite of impaired FFA utilization, further corroborates systemic energy deficiency. These findings collectively emphasize the non-redundant role of glucose in neonatal energy homeostasis, particularly under conditions of cytoskeletal dysregulation.

Our investigation into hepatic gluconeogenesis revealed that *Fscn1* KO did not impede the canonical hormone-regulated transcriptional gluconeogenic pathway, thereby excluding hyperinsulinemic hypoglycemia as a potential etiology. Circulating insulin concentrations and downstream insulin signaling target — AKT phosphorylation status remained unaltered in *Fscn1* KO neonates. Transcriptomic analysis revealed selective disruption of glycerol-mediated gluconeogenesis in KO mice, notably Gpd1 and Gpd2, which is an alternative metabolic route governed by substrate availability and intracellular redox homeostasis [24]. Systematic validation across *in vitro* and *in vivo* models established that *Fscn1* deficiency causes significant downregulation of GPD1 expression, perturbs mitochondrial and cytoplasmic [NADH:NAD^+^] redox equilibrium, and reduces hepatic glycerol availability. The functional deficit in glycerol-to-glucose conversion was further confirmed by impaired glycemic recovery in *Fscn1* KO mice during glycerol-tolerance tests. Intriguingly, while both *Fscn1* KO mice and derived cell lines exhibited reduced Gpd1 mRNA and protein expression, the regulatory interplay FSCN1 and GPD1 appears indirect for their distinct subcellular localization.

FSCN1 has been considered as a marker of immature hepatocyte [55], its level progressively decreases during early liver development [56]. Notably, the impaired hepatic glyceroneogenesis in *Fscn1* KO mice was not associated with hepatocyte injuries. While perivascular nuclear aggregation of hepatocytes adjacent to portal veins, a histological feature suggestive of subclinical injury [57], was detected in *Fscn1* KO neonates. Quantitative assessment of liver function enzyme levels including alanine aminotransferase (ALT), aspartate transaminase (AST) and AST/ALT ratio (Fig. S2E-F) revealed no evidence of substantial hepatic damage. This collectively reinforces the notion that *Fscn1* KO induces hypoglycemia through a selective disruption of the glycerol-gluconeogenic pathway rather than through impairment of the hepatocytes structure. This study highlights FSCN1’s critical role in early hepatic gluconeogenesis, suggesting that the metabolic regulation during liver development has been underestimated, which might provide a clinical reference for metabolic abnormalities caused by cytoskeletal deletions.

In conclusion, we redefine FSCN1 as a linchpin connecting actin dynamics to neonatal energy homeostasis. By orchestrating the glycerol gluconeogenic shunt, FSCN1 ensures glucose output during the critical postnatal transition, offering novel insights into congenital hypoglycemia and cytoskeletal-metabolic crosstalk. These findings pave the way for exploring cytoskeletal targets in metabolic disorders and refining neonatal hypoglycemia management strategies.

## Acknowledgements

We thank Dr. Jing Li for their helpful suggestion and comments on the manuscript; Dr. Chunming Guo for supporting pathological analysis platform.

## Supplementary Data

**Figure S1.**
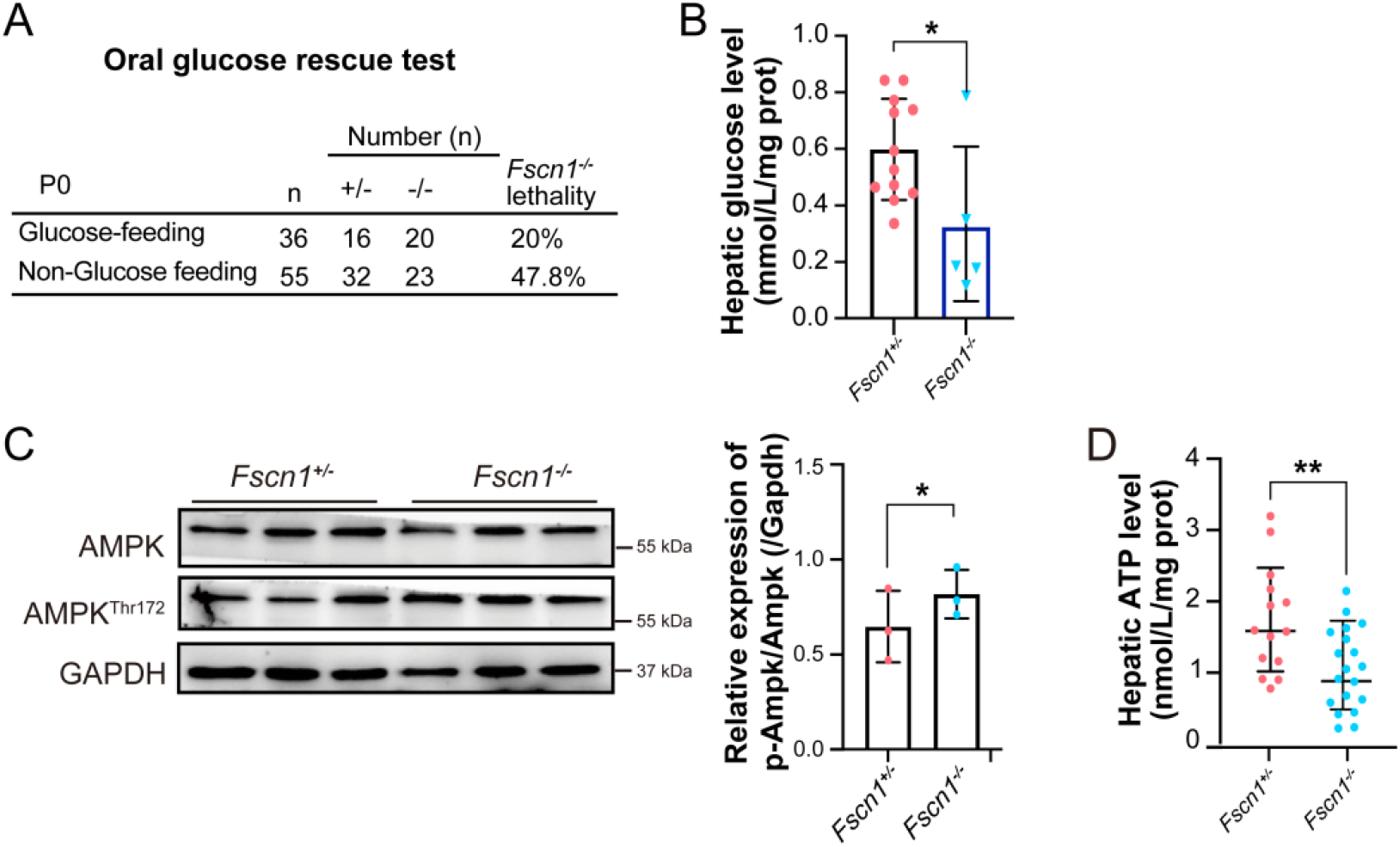
Metabolic consequences of *Fscn1* deficiency in neonatal mice. (A) Oral glucose rescue test in P0 *Fscn1*^*+/-*^ and *Fscn1*^*-/-*^ mice, with lethality rates of *Fscn1*^*-/-*^ mice were recorded. (B) Hepatic glucose levels in *Fscn1*^*+/-*^ and *Fscn1*^*-/-*^ P0 mice. (*Fscn1*^*+/-*^, n=12; *Fscn1*^*-/-*^, n=5). (C) Western blot analysis of AMPK activation in *Fscn1*^*+/-*^ and *Fscn1*^*-/-*^ P0 mice liver, and qualified using Image J (each group, n=3). (D) Hepatic ATP levels measured in *Fscn1*^*+/-*^ and *Fscn1*^*-/-*^ P0 mice. (*Fscn1*^*+/-*^, n=14; *Fscn1*^*-/-*^, n=19). All mice used in these experiments were littermates. Data are presented as mean ± SEM, statistical significance was determined by two-tailed t-test, with *\*p* < 0.05, *\*\*p* < 0.01,

**Figure S2.**
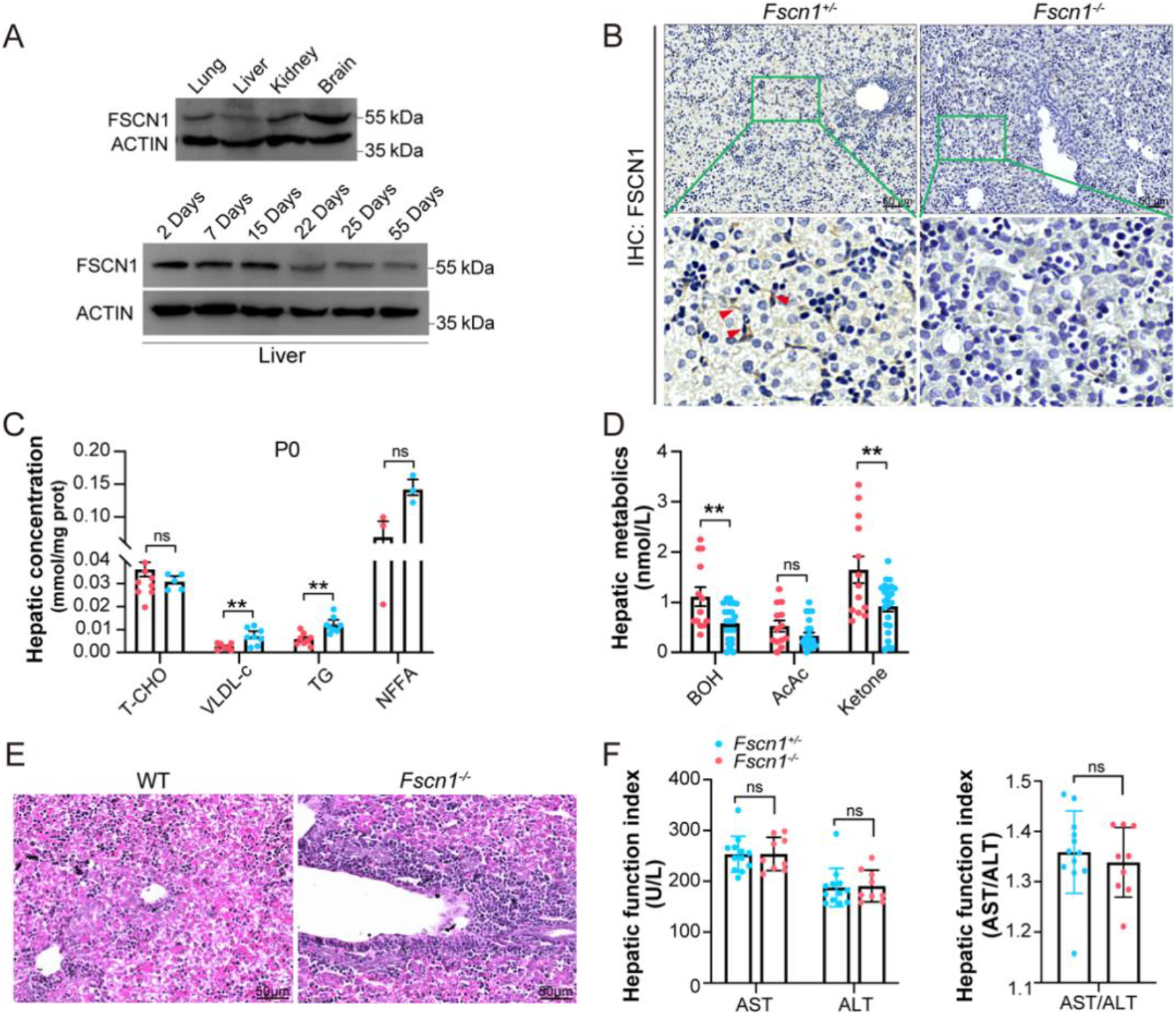
Tissue-specific FSCN1 expression and metabolic effects of *Fscn1* deficiency in neonatal mice. (A) Protein expression levels of FSCN1 in lung, liver, kidney, and brain tissues in WT mice at P14, and in liver tissues from mice aged 2 to 55 days. (B) Immunohistochemistry of FSCN1 in P0 neonatal mouse liver slices. Scale bar=50 μm. Red triangles indicate FSCN1-positive staining (each group, n=3). (C) Hepatic lipid levels in P0 *Fscn1*^*+/-*^ and *Fscn1*^*-/-*^ mice, including total cholesterol (T-CHO), very low-density lipoproteins (VLDL-c), triglycerides (TG) and non-esterified fatty acid (NEFA) and the lipids contents were quantified by BCA method. For T-CHO, VLDL-c and TG, each group included≥5 mice; for NEFA, n=3 per group. (D) Hepatic levels of β-hydroxybutyrate (BOH) acetoacetate (AcAc) and total ketone in *Fscn1*^*+/-*^ and *Fscn1*^*-/-*^ P0 mice (each group, n≥13). (E) Hematoxylin and eosin (H&E) staining of P0 mouse liver tissue; scale bar=50 μm (each group, n=3). (F) Levels of Serum liver function markers Alanine aminotransferase (ALT), Aspartate aminotransferase (AST) in *Fscn1*^*+/-*^ and *Fscn1*^*-/-*^ mice and their ratio (AST/ALT). (*Fscn1*^*+/-*^, n=12; *Fscn1*^*-/-*^, n=9). All mice in these experiments were littermates. Data are presented as mean ± SEM, statistical significance was determined by two-tailed t-test, with *\*p* < 0.05, *\*\*\*p* < 0.001.

**Table S1.**
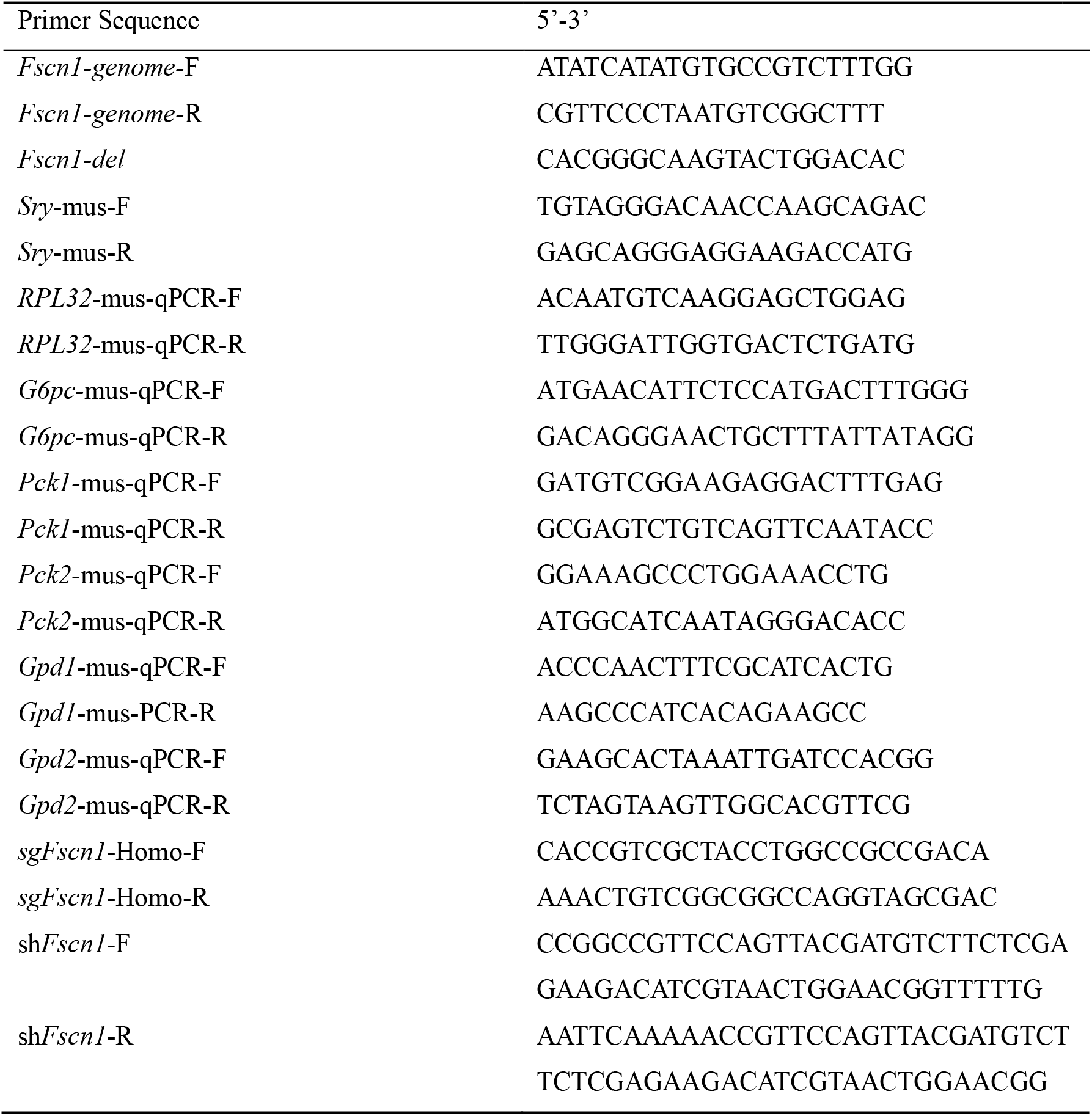
Primers used in this study.

**Table S2.** Reagents used in this study.

| Reagents | Source | City | Country |
| --- | --- | --- | --- |
| Alanine transaminase (ALT) | Nanjing Jiancheng Bio | Nanjing | China |
| Aspartate transaminase (AST) | Nanjing Jiancheng Bio | Nanjing | China |
| ATP Assay | Nanjing Jiancheng Bio | Nanjing | China |
| Nonesterified Free fatty acid (NEFA) | Nanjing Jiancheng Biog | Nanjing | China |
| Triglyceride (TG) | Nanjing Jiancheng Bio | Nanjing | China |
| Total Cholesterol (T-CHO) | Nanjing Jiancheng Bio | Nanjing | China |
| Glycerol | Nanjing Jiancheng Bio | Nanjing | China |
| Glucose | Nanjing Jiancheng Bio | Nanjing | China |
| High-density lipoprotein cholesterol (HDL-C) | Nanjing Jiancheng Bio | Nanjing | China |
| Very Low-density lipoprotein cholesterol (VLDL-c) | Nanjing Jiancheng Bio | Nanjing | China |
| Ketone body Assay | Solarbio | Beijing | China |
| Insulin Elisa Kit | Jonln Biotechnology | Shanghai | China |
| D-glucose | Sigma | St. Louis | USA |
| Protease inhibitor cocktail | MCE | Shanghai | USA |
| Glycerol | Sangon Biotech | Shanghai | China |
| TriZol | Lablead | Beijing | China |

**Table S3.**
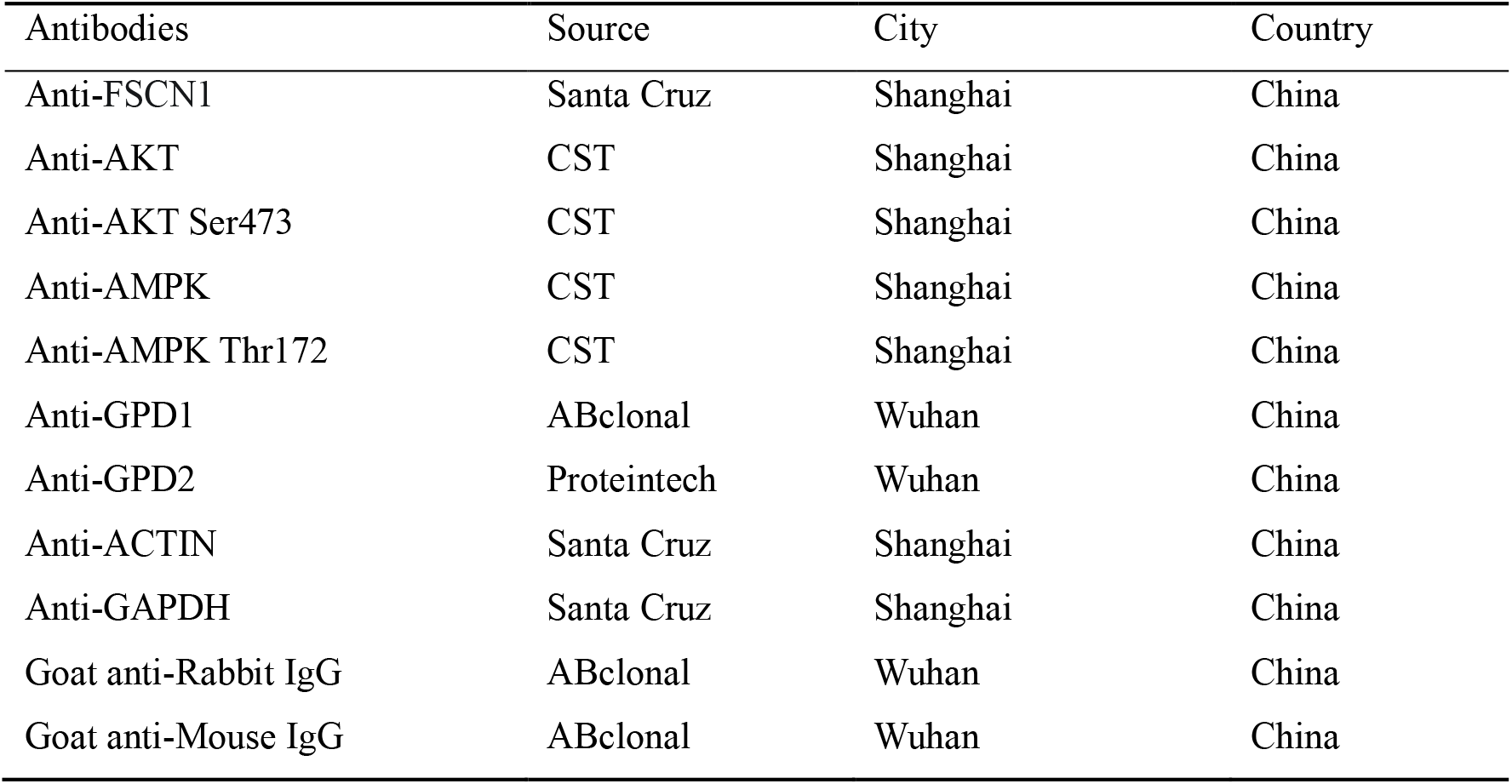
Antibodies used in this study.

